# A dependency-free, streamable format and cross-language toolkit for scalable LC-MS feature detection: reading only what you need

**DOI:** 10.64898/2026.09.15.751169

**Authors:** Javier Osorio Mosquera, Nathan G Lawler, Maartje Cox, Jorge Osorio Mosquera, Wargner A Moreno L, Samuele Sala, Vimalnath Nambiar, Luke Whiley, Jeremy K Nicholson, Elaine Holmes, Julien Wist

**Affiliations:** Centre for Computational and Systems Medicine, Health Futures Institute, Harry Perkins Institute of Medical Research, Murdoch University, Perth, Western Australia 6150, Australia; School of Allied Health (Exercise Science), College of Health and Education, Murdoch University, Perth, Western Australia 6150, Australia; School of Food Engineering, Universidad del Valle, Melendez 76001, Cali, Colombia; Chemistry Department, Universidad del Valle, Melendez 76001, Cali, Colombia; Curtin Medical Research Institute, Curtin University, Bentley, Perth, Australia; School of Diagnostic and Therapeutic Sciences, Curtin University, Bentley, Perth, Australia; Dementia Centre of Excellence Enable Institute, Curtin University, Bentley, Perth, Australia; School of Medicine, The Hong Kong University of Science and Technology, Clear Water Bay, Kowloon, Hong Kong; Department of Metabolism, Digestion and Reproduction, Faculty of Medicine, Imperial College London, Sir Alexander Fleming Building, South Kensington, London SW7 2AZ, UK

## Abstract

Mass spectrometry generates data faster than it can be read, and the exchange standard, mzML, is text-based and must be parsed in full before any spectrum is accessible. Binary alternatives are smaller but depend on storage engines such as HDF5, so access is dictated by the engine, not the file. We present Ionic (.ion), an open-source, compact, streamable binary format, and Quant·ion, a processing toolkit built on the former. Ionic stores spectra, chromatograms and metadata as independently compressed, indexed blocks, so a reader retrieves only the bytes it needs, even inside a web browser, and converts losslessly to and from mzML. Ionic was smaller than compressed mzMLb on every acquisition type tested, and extracting one compound took under 40 ms, 35 to 90 times faster than an mzML reader. Quant·ion exposes one core to R, Python, and JavaScript with identical results, and recovered 97% of true features on a ground-truth benchmark.

**Graphical Abstract:** 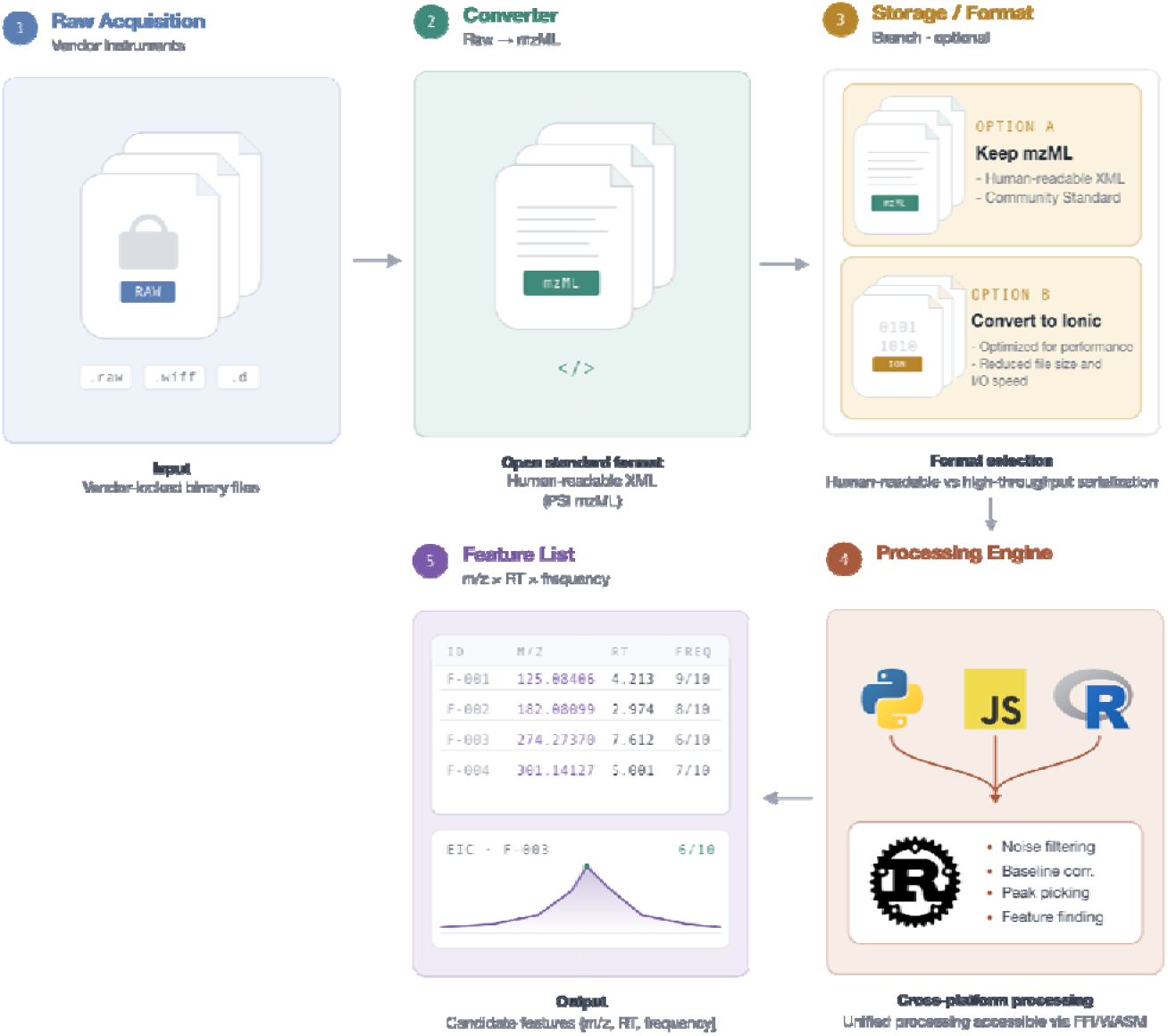

## Introduction

Mass spectrometry has become one of the principal sources of large-scale data in the life sciences, including metabolic phenotyping, proteomics, and imaging. As studies grow, the computational bottleneck is shifting from statistical modelling to the sheer cost of moving and reading data. Instruments now generate data faster than most laboratories can store, move, and analyze. Disk space is the smallest part of that burden. Once a study reaches the terabyte scale, analysis moves off the researcher’s own machine and onto dedicated servers. Every processing pass over the data becomes a job to schedule. For example, liquid chromatography - mass spectrometry (LC-MS) files have grown with every gain in mass resolution and scan speed, with recent platforms even adding an additional dimension. Ion mobility spectrometry inserts collision cross-section as a fourth measurement alongside retention time, m/z ratio, and intensity. Similarly, mass spectrometry imaging records a full spectrum at every spatial position, so a single tissue section can reach tens of gigabytes. The data swells with each new innovation so that a single injection is acquired faster than it is read back.

Contemporary cohorts comprise thousands of acquisitions collected across multiple instruments, laboratories, and research centres, and these are increasingly harmonised and fused with other omics layers before being used to train machine learning models. At population scale, that dataset far exceeds available memory, so that every training iteration is dominated by data access time. Making the data more compact and faster to read from storage therefore substantially accelerates learning procedures while reducing computational and energy requirements.

Scale creates a second problem. Modern platforms report thousands of features per sample, but the underlying peaks are rarely examined, either by the analysts who generate these, or by peers reviewing the resulting claims, because the files are large and the tools required to open them are cumbersome to install and operate^18^. The two problems reinforce each other: machine learning depends on carefully curated features and yields thousands of new candidates that themselves require review. If any individual feature could readily be retrieved from a remote file and displayed in a web browser, inspection would become a routine part of analysis rather than an exceptional effort, making the evidence behind clinical and biological conclusions auditable. Recognising that reproducibility matters as much as detection accuracy, recent tools have begun to address provenance and feature correspondence explicitly ^1^.

Prevailing data formats work against both objectives. The community standard, mzML, is a text-based format ^2^ chosen for its readability. To keep final file size in check, the numerical arrays that dominate its size are routinely encoded and compressed. As a result, readability is mostly lost. Binary formats recover much of the lost space by building on general-purpose storage engines (mzMLb, ^3^; mzDB, ^4^; mz5 ^5^). This choice of technology ties every downstream data consumer to a specific software dependency, adding a layer of complexity in the lightweight environments where inspection could be most efficient (i.e. a web browser). Recent work has proposed adopting established columnar formats, confirming that a compact, binary, streamable representation is the right objective, while inheriting the same constraint ^6^. To compound to this issue, most instruments write proprietary formats that must first be converted to an open standard ^7^. This lack of a widely adopted, efficient file format forces laboratories to either retain both the original and the converted data files, inflating archives and complicating long-term data management, or to add a computationally expensive and time-consuming step to their pipeline.

Processing software is similarly constrained. Many tools remain bound to a particular operating system, graphical interface, or programming ecosystem, restricting where analysis can take place (MS-DIAL, ^8^; and the vendor packages MarkerView and Compound Discoverer, described in Methods Section). Virtualisation technology mitigates this at the cost of additional setup, maintenance, and computational overhead ^9^. In multidisciplinary research teams, where expertise is distributed across medicine, biology, analytical chemistry, statistics, and computer science, the requirement for a specific programming environment is itself a barrier to adoption and to independent verification ^1,10,11^. Furthermore, scaling compounds the difficulty, because optimal processing parameters depend on the dataset, so that large studies are commonly processed multiple times. Maintaining and versioning such a fragmented code base and pipeline is time-consuming and prone to errors.

Here we present Ionic, an open and streamable binary format for mass spectrometry data, and Quant·ion, a processing toolkit built upon it. Ionic stores spectra, chromatograms, and metadata as independently compressed blocks indexed by compact directories, so that any part of a file can be retrieved without reading or decompressing the rest. Conversion to and from mzML is lossless, so it fits into existing data pipelines. Since the format depends on nothing beyond ordinary file access and a standardised decompressor, a complete reader can be implemented in a few hundred lines of code in essentially any language, including one that runs inside a web browser. Quant·ion processes very large files at low and near-constant memory, and exposes a single code implementation to R, Python, and JavaScript, etc., so that the same acquisition returns identical results in every programming environment. Finally, we validated our workflow by applying these tools across all study cohorts to build a unified, accessible database for both our targeted and untargeted assays.

## Results

### Design of the Ionic file format

Ionic (.ion) is a little-endian binary format that holds the same content as mzML, spectra, chromatograms and metadata. Conversion to and from mzML is lossless. Ionic is structured so that a program can work with very large files by reading only the chunks it needs for a given query, leaving the rest compressed (Figure 1). Two requirements follow from that objective, shaping the whole design; a reader must be able to locate any part of the file without scanning it, and be able to decode that data without an external storage engine.

**Figure 1.**
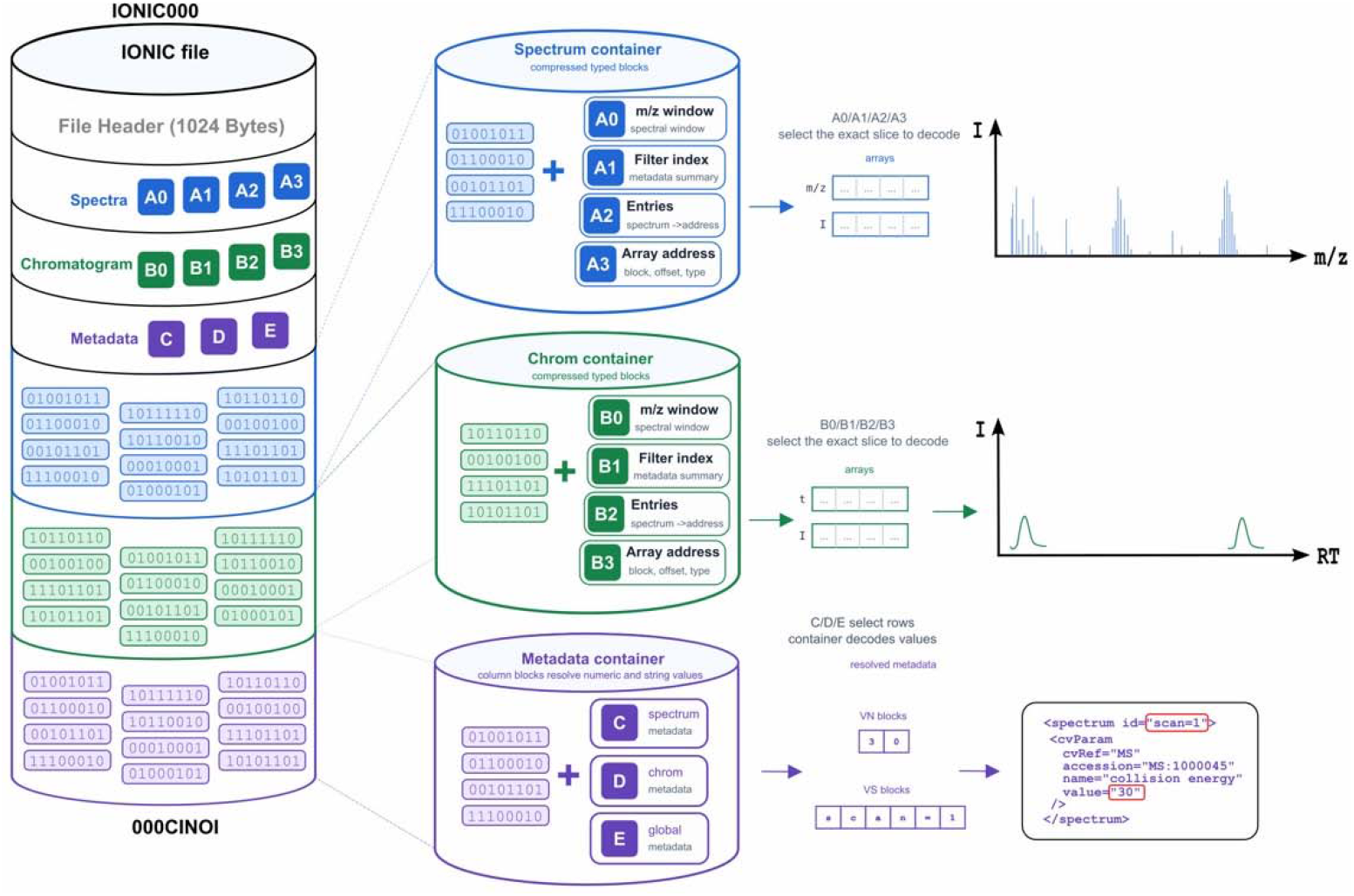
Architecture of the Ionic binary format. An Ionic file is organised into three independent containers: spectra (blue, sections A0 - A3 and the spectrum block container), chromatograms (green, sections B0 - B3 and the chromatogram block container) and metadata (purple, sections C - E). All section offsets, item counts, and CRC-32 integrity checksums are stored in a fixed 1024-byte file header. A fixed 8-byte trailer written last marks the end of the file. Within each data container, four compressed index sections (window directory, filter summary, array index, address table) allow the decoder to resolve a query to a list of blocks and decompress exactly those, without reading the whole file. Metadata sections use a columnar parallel-array scheme that resolves PSI-MS CV (Proteomics Standards Initiative Mass Spectrometry controlled vocabulary) parameters to human-readable values at runtime (see Supplementary Material S1).

Every Ionic file begins with a fixed 1024-byte header recording the byte offset and length of every section, the number of spectra and chromatograms, per-section CRC-32 integrity checksums, and the global encoding flags. Because the header is fixed in size and position, a parser reads it in a single request and computes the position of any section arithmetically. A fixed 8-byte trailer, written last, and the opening signature together confirm that the file is not truncated. The signature, trailer and all section checksums are verified prior to any data access.

Array blocks are addressed via four index sections per container: an m/z-window directory (A0), a fast filter summary holding the scalar fields used to filter spectra (A1), a spectrum array index (A2), and an array address table giving the block, element offset and element count of every array segment (A3); chromatograms use the same arrangement through sections B0 - B3. These sections are compressed on disk and decompressed when the file is opened, and their records are fixed-width, so any record is located by computation rather than by scanning. A query therefore resolves to a list of block identifiers before a single byte of signal is decompressed, while remaining blocks stay compressed and are never loaded.

Encoding follows the behaviour of the values rather than a generic rule. All numeric arrays are stored as native types (16-, 32- and 64-bit integers and floats) and packed into independently compressed blocks. The uncompressed size of blocks is configurable and is recorded in the header. *m/z* ratio, time and ion-mobility float arrays are delta-coded, exploiting their monotonicity, while intensity and integer arrays are stored as-is. Each block is then byte-shuffled and compressed with Zstandard. Metadata is held separately in sections C, D and E as compressed columnar arrays of controlled-vocabulary (CV) parameters. These are grouped so that a reader decodes only the group it needs. Human-readable names are never stored: the CV prefix becomes a 1-byte code and the accession a 32-bit unsigned integer, resolved at read time against a standard CV table.

As a result, reading an Ionic file only requires byte-level file access and a Zstandard decoder. While HDF5, SQLite and Apache Parquet own the structure of the file and determine how data can be accessed, Zstandard [17] is only used for block compression. Their positions, lengths and contents are described by the header and the index sections. A complete reader is therefore a few hundred lines of code in any language, including one that compiles to WebAssembly and runs inside a web browser. We make further use of this property later in the paper. The full specification is given in Supplementary Material S1.

### Storage Footprint of the Ionic format

Since Ionic is designed for mass spectrometry data rather than built on a general-purpose container, it can encode each array according to how its values behave, so the savings will depend on the acquisition type rather than the file size (Figure 2). In dense profile acquisitions the gain comes from delta-coding and byte-shuffling the monotonic m/z axis, whereas in centroided or targeted acquisitions that redundancy is largely absent and the gain comes instead from replacing the ASCII text and XML overhead with native numeric types and columnar metadata. Across the six acquisition types tested (Figure 2), Ionic files were ∼1.3× to ∼17.7× smaller than mzML: ∼17.7× for the targeted lipidomics assay (SCIEX 6500 and 7500), and ∼2.5× to ∼2.9× for the semi-targeted, untargeted (Bruker Impact II) and DDA proteomics assays (See Methods for details). As expected, the advantage over mzMLb was narrower and more uniform, ∼1.07× to ∼1.59×, which is still substantial for large datasets. The imaging dataset, an imzML acquisition, is stored as two files: the .imzML metadata and the .ibd binary data, while Ionic stores it in a single file that is still ∼1.3× smaller than the two combined.

**Figure 2.**
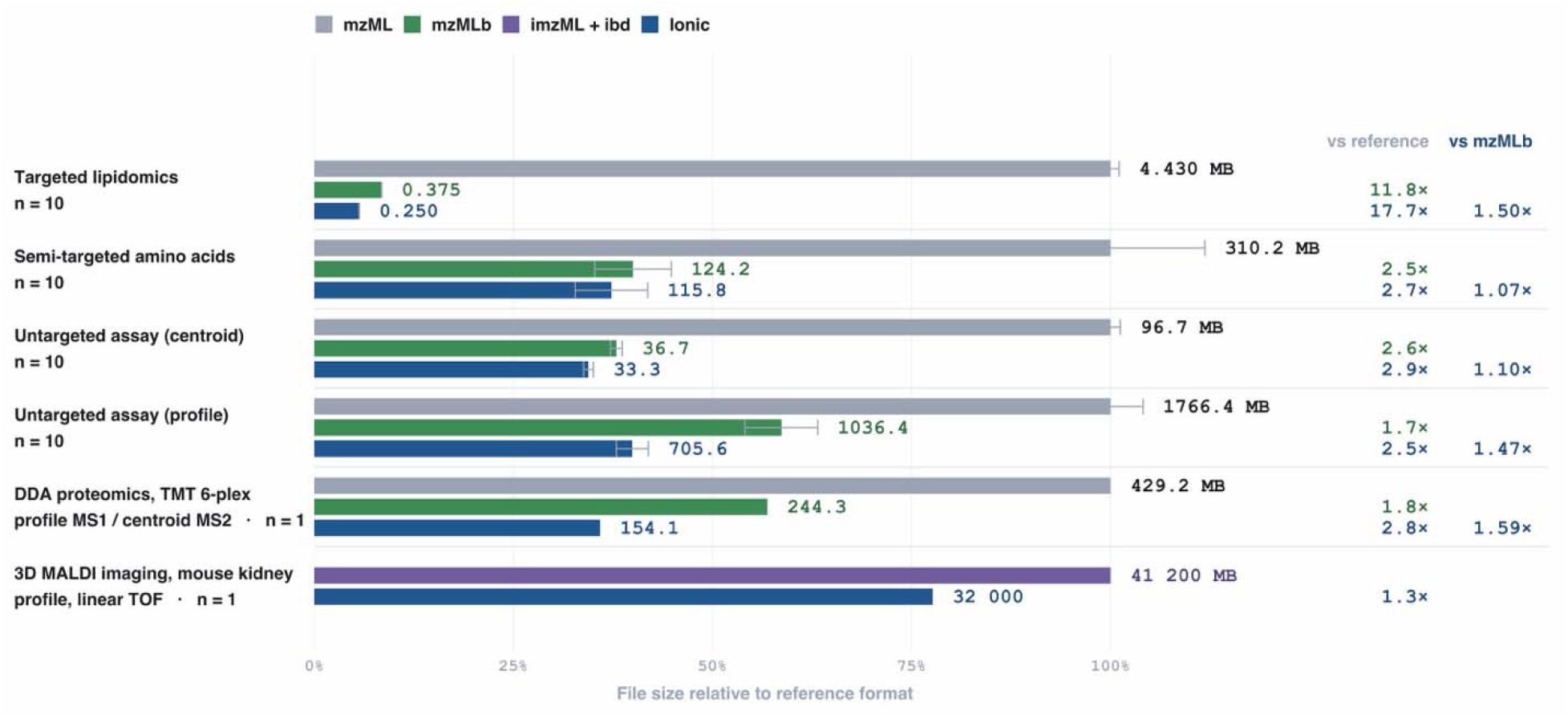
File size across formats for six acquisition types. mzML, mzMLb and Ionic (.ion) file sizes for a targeted lipid assay, a semi-targeted amino-acid assay, centroided and profile-mode untargeted acquisitions, a DDA TMT 6-plex proteomics run (profile MS1, centroid MS2), and a 3D MALDI imaging dataset of mouse kidney (profile, linear TOF). The bar length indicates the file size relative to mzML within each acquisition type. Absolute sizes in MB and number of files are displayed for each bar. Bars show the mean and error bars indicate ± 1 s.d. across all files. No error bar is shown where n < 5. Grey values give the fold reduction versus mzML, blue values in parentheses the fold reduction of Ionic versus mzMLb. imzML is already a binary format. Proteomics data were accessed from PRIDE PXD00000 ^12^; MALDI imaging data from Oetjen et al. ^13^.

For large research institutes (i.e. phenome centres) the effects can be seen to compound. At the time of writing, 103,195 acquisitions from six assays, roughly 4.5 TB (in compressed_vendor_format.tar.xz), have been converted to Ionic and form a single streamable archive (Supplementary Table S7).

### A cross-language toolkit built on Ionic

Quant·ion provides peak picking, peak fitting, baseline correction, noise-level estimation, extracted-ion chromatogram (EIC) calculation and feature detection, and accepts both .ion and .mzML input. The core is written in Rust and exposed to R, Python and JavaScript, so every language calls the same code and returns exactly the same answer. This core compiles to WebAssembly and thus can be executed inside a web browser on streamed blocks. The result is also independent of the input format. For one test compound in each of the three datasets, the EIC maximum agreed to all reported digits between Quant·ion reading Ionic and independent readers of mzML and mzMLb (Supplementary Table S6).

### Streaming access and in-browser inspection

Partial reads are a property of the format, while turning them into remote processing is the job of the toolkit. The file does not need to be present on the local machine: a URL to a remote file is enough. Quant·ion reads the small header, maps the file layout, and issues HTTP range requests only for the blocks required (Figure 3). Fetched blocks enter a fixed-capacity cache, so memory usage is determined by the cache’s capacity rather than by file size, i.e., a multi-gigabyte acquisition on a remote server can be queried as if it were local. Inspecting a single feature across multiple samples, for example overlaying their extracted-ion chromatograms to confirm a peak visually, transfers and decompresses only a few blocks from each file.

**Figure 3.**
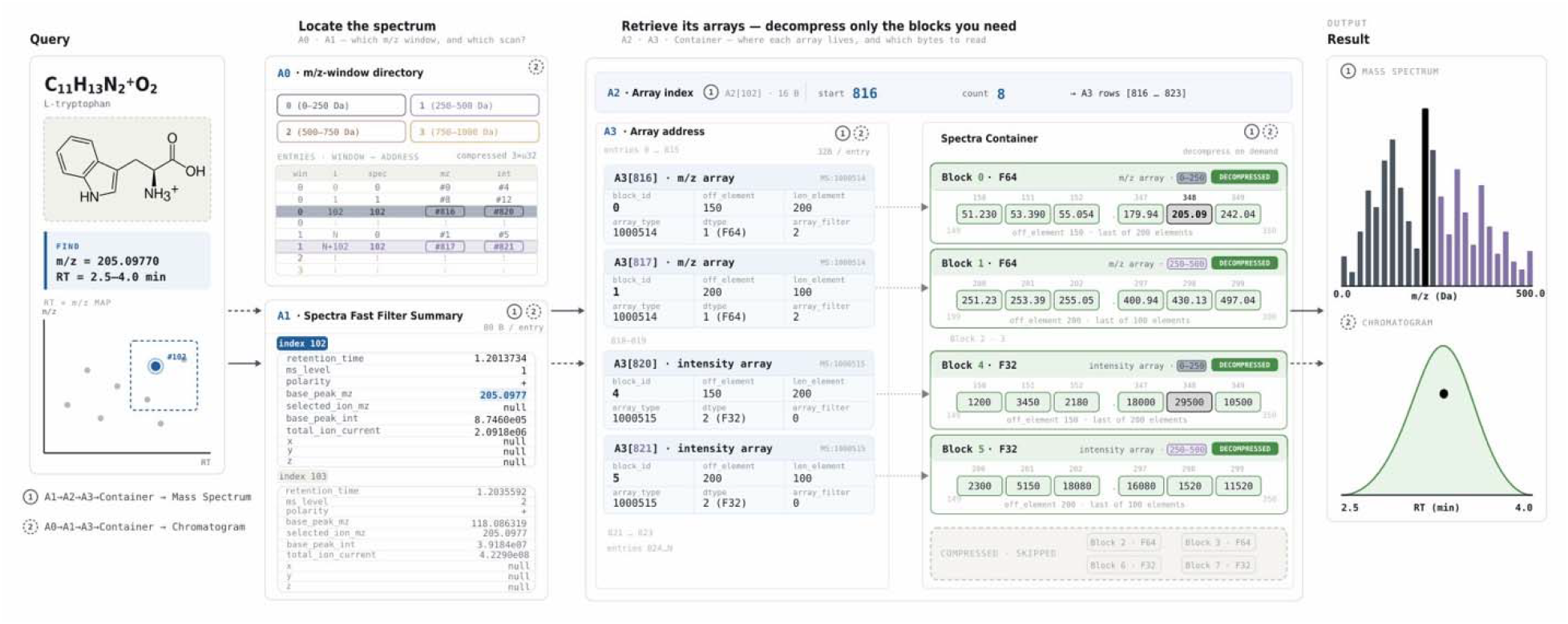
Locating and reading one spectrum. Spectrum data is organised across four index sections, A0 (m/z-window directory), A1 (fast filter summary), A2 (spectrum array index) and A3 (array address table) and one compressed spectrum container. The worked example resolves a query for L-tryptophan ([M+H]□, *m/z* 205.09715, retention time 2.5 - 4.0 min) to spectrum index i = 102.

Because a query is resolved from the index before any signal is decompressed, the time required for opening a file is largely decoupled from its size. Opening and extracting one compound took 12 to 35 ms with Quant·ion on Ionic, against 0.5 to 3.2 s for an mzML reader on the same acquisitions (Supplementary Table S6). Since no single implementation reads all three formats, these comparisons require different readers. The mzMLb timings were therefore obtained through a Python composite reader, so the results reflect both the reader and the format, which severely limits the value of this comparison. We did not implement mzMLb reading in Quant·ion, because doing so would mean carrying an HDF5 runtime into every target we support, including WebAssembly. The comparison with mzML does not carry this ambiguity: Quant·ion reads both formats, and with the same code on both, EIC extraction from Ionic was 10 to 120 times faster than from the source mzML across every encoder configuration tested (Figure 4).

**Figure 4.**
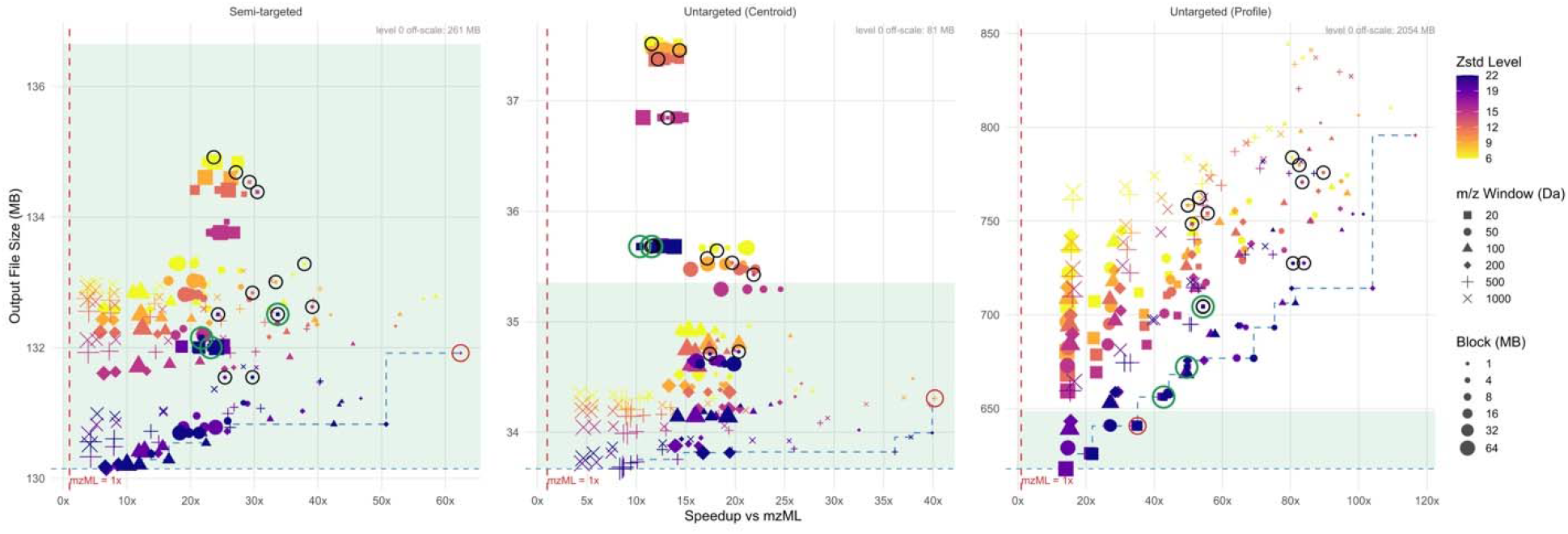
File size and EIC calculation speed for different Ionic settings. Panels are three files, one from each of three LC-MS datasets: semi-targeted, untargeted centroid and untargeted profile. The reference is the EIC panel computed from the source .mzML with Quant·ion, the red dashed line is that 1× reference, annotated with the .mzML file. All other markers are Ionic files, read with the Ionic Python wrapper (v0.1.0). Each marker is one encoder configuration (216 per dataset: 6 zstd levels × 6 block sizes × 6 m/z windows), colour gives the zstd compression level, shape the m/z window, area the block size. The green band spans sizes within 5% of the smallest .ion (blue dashed line), the blue step traces the best trade-off.

The defined size of the blocks and the window width affect the final size of the encoded file, since Zstandard compresses larger blocks better. In turn, smaller blocks mean a smaller cache and more granular access to the features. Figure 4 extends this across 216 encoder configurations on three acquisition types: the size penalty stays within a few percent for semi-targeted and centroided data but reaches a third for profile data, where block size and window width must be traded against granularity, while the speed advantage over mzML holds at every configuration. Extracting alanine with 1 MB blocks and a 20 Da window transfers 4.4 MB, 6.5% of the file, in 0.17 s. The same query using 64 MB blocks and a 500 Da window transfers 57.0 MB, 85%, in 16.2 s. The cost per metabolite falls when many are read from a single file; with a 500 Da window, several metabolites are served by the same blocks, whereas with a 20 Da window multiple blocks need to be fetched separately. Large blocks therefore suit untargeted screening, small blocks targeted assays.

As a demonstration, we deploy a web application [https://phenological.github.io/ion-beam/]. It opens an Ionic file directly from a URL and lets the user browse spectra, select a feature and display its extracted-ion chromatogram, in the browser, on a laptop, with no installation. It is a static webpage (a standalone .html file) that keeps working as long as the archived file is reachable. A collaborator, reviewer or curator can therefore re-examine the raw evidence behind any reported feature from a single link, contributing towards the reproducibility and provenance goals that motivated the format.

### Quant·ion + Ionic, evaluation on Untargeted LC-MS benchmark

To test the format and the toolkit together on acquired mass spectrometry data, we processed the standard-mixture benchmark used in the study of Li et al (2018), where two mixtures were measured containing 1,100 drugs and metabolites, 130 of which were spiked at defined SB:SA ratios to simulate biomarkers. The mixtures were acquired using both a Sciex TripleTOF 6600 and a Thermo Q Exactive HF (QE HF) where 970 (Sciex) and 836 (Thermo) features were annotated and confirmed. The 68 and 50 spiked compounds that met the source study’s marker criteria serve as ground truth for discriminating markers. Comparator accuracy values are taken from Li et al. (2018) as published. Using Quant·ion we recovered 943 of 970 true features (97.2%) on the Sciex TripleTOF 6600 and 808 of 836 (96.7%) on the QE HF (Figure 5a). Of the 970 entries in the TripleTOF 6600 list, 89 items are reported with the same m/z and retention time than at least one other item (44 pairs), 2 other items (43 triads), and 3 other items (1 quartet). Since they are isobaric and reported as co-eluting, a single peak is expected. However, closer inspection shows that those peaks are well resolved (Supplementary Note 2). We detected a feature at 43 of these 44 positions and counted every compound there as found. This increased our recovery from 899 of 970 (92.7%) to 943 (97.2%).

**Figure 5.**
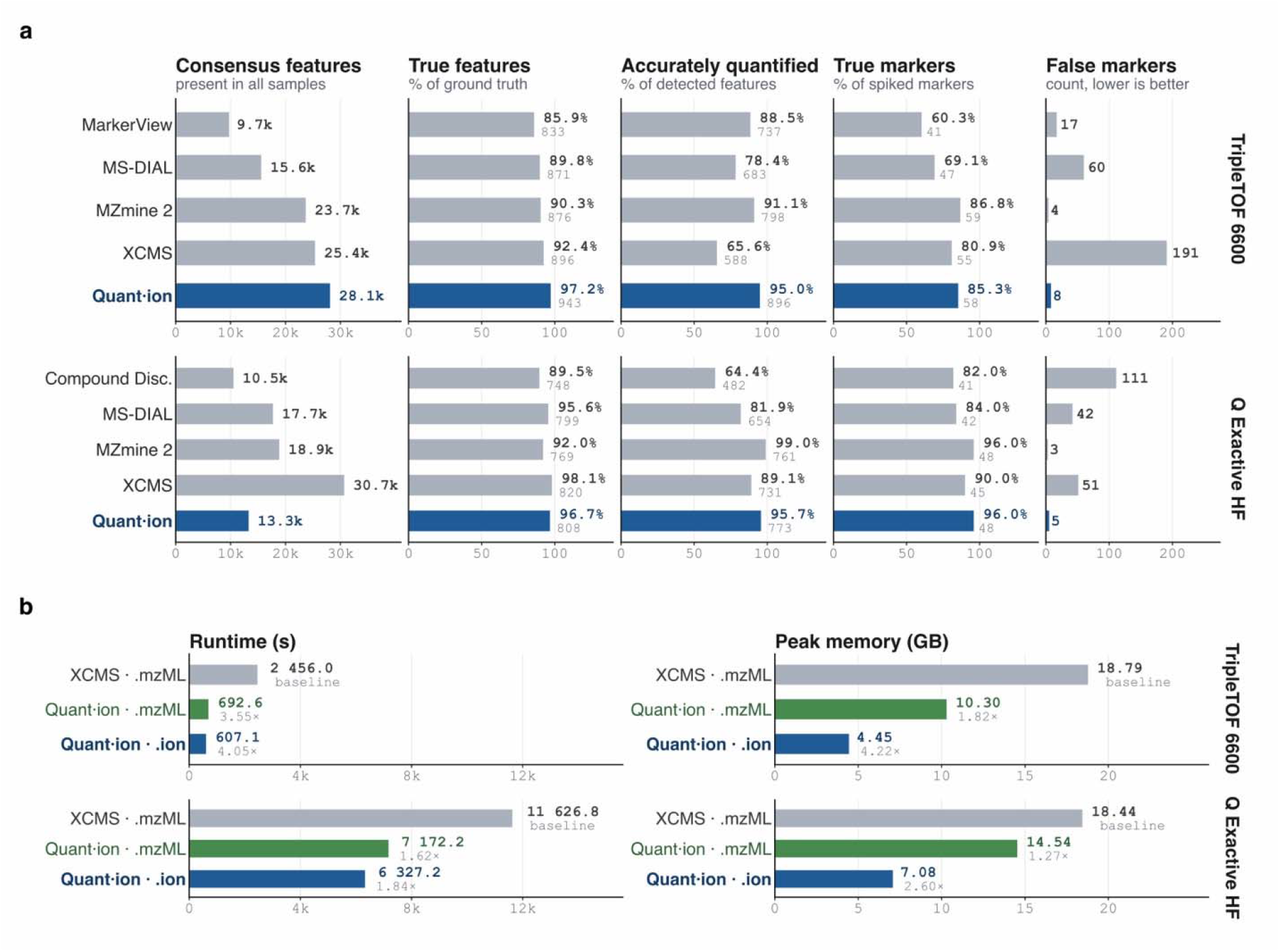
Feature detection and processing cost on a ground-truth benchmark. Standard mixtures SA and SB acquired on a SCIEX TripleTOF 6600 and a Thermo Q Exactive HF (QE HF), with 970 and 836 true features and 68 and 50 spiked markers as ground truth. **a.**Detection and quantification per software. “Consensus features” are detected in every sample. “Accurately quantified” means the SB:SA fold change matches within 20%, and is conditional on detection. “False markers” are false positives. Comparator values are taken from Li et al. (2018) **b**. Runtime and peak memory. Bars are relative to the XCMS baseline achieved in-house. Grey values give the fold reduction versus that baseline. Blue values on the .ion row give .ion versus .mzML for the same tool. Quant·ion was called through its R wrapper (v0.1.0). CPU only was allowed with the same thread count for both tools (Methods). All runs were performed on a 2019 Mac Pro (3.5 GHz 8-core Intel Xeon W, 32 GB).

Of the true features it detects, Quant·ion recovers the benchmark expected fold change within 20% variation for 896 of 943 (95.0%) on the TripleTOF 6600 and 773 of 808 (95.7%) on the QE HF, consistent across both platforms. The comparator rates are lower and more variable: 65.6% (XCMS) to 91.1% (MZmine 2) on the TripleTOF 6600, and 64.4% (Compound Discoverer) to 99.0% (MZmine 2) on the QE HF. Since this metric is conditional on detection, a tool that recovers fewer features is scored on a smaller and often easier set.

Both detection and quantification combine for marker recovery. Quant·ion flags 58 of 68 simulated biomarkers (by spiking) (85.3%) on the TripleTOF 6600 with 8 false markers, and 48 of 50 (96.0%) on the QE HF with 5. Only MZmine 2 reports fewer false markers on both platforms (4 and 3); XCMS (191, TripleTOF 6600) and Compound Discoverer (111, QE HF) report by far the most. Only Quant·ion and MZmine 2 combine high recovery, accurate quantification and a low false-marker count on both platforms.

We then compare processing costs against XCMS^10^, the standard and established open-source tool, since the other software runs through a graphical interface or on a single operating system, precluding a controlled measurement. The same workstation (see Online methods section 4.4) was used, and the achieved accuracy is comparable: 943 versus 934 of 970 true features on the TripleTOF 6600, and 808 versus 805 of 836 on the QE HF.

In contrast, figure 5b shows that Quant·ion reads and processes TripleTOF 6600 data 4× faster (607.1 s) than XCMS (2,456.0 s) and uses 4× less memory at runtime (4.45 GB against 18.79 GB). For QE HF data the improvement is a 1.84-fold speed up and 2.60-fold reduction in memory, from 6,327.2 s with 7.08 GB for Quant·ion against 11,626.8 s and 18.44 GB for XCMS. The QE HF dataset has a greater number of MS1 spectra, and more points per spectrum, accounting for the longer runtime.

## Discussion

Untargeted LC-MS feature detection has long forced a choice between speed, memory efficiency, and accessibility. Our results demonstrate that all three can be achieved in tandem. Ionic files were smaller than compressed mzMLb across every acquisition type tested (∼1.07× to ∼1.59×), with lossless conversion in both directions. Quant·ion matches or exceeds the true-feature recovery of established tools on a benchmark dataset with a defined chemical ground truth. It delivers one of the lowest false-marker counts on both platforms, and executes 1.8-to 4.0-fold faster than XCMS using 2.6-to 4.2-fold less memory. The toolkit runs natively on Windows, macOS, and Linux and can be executed within R, Python, and JavaScript, returning identical results for each.

A general-purpose storage engine such as HDF5 controls the structure of the file, and performance is dependent on the available implementations for that engine in any given language and platform. By contrast, reading an Ionic file only requires byte-level file access and a Zstandard decoder. The layout of the blocks is controlled entirely by the header and the index sections. This design allows for per-feature inspection inside a browser tab. A static web application that opens an archived file from a URL with no installation can be deposited along with the data to ensure permanent visibility, facilitating the curation, review and re-use of the peaks and features that support biomarker discovery.

Ionic is not the first attempt to replace mzML with a compact binary representation, and mzPeak [5] shares our view that a compact, binary, streamable format is the right objective. The two proposals differ in that our design avoids inheriting a data engine and the cost it presupposes at runtime. Ionic inherits nothing and therefore implements only what it needs. We are also wary of betting on a community-wide vendor alignment; this has not happened in the two decades since mzML. For real time processing, the “last mile” requires a solution to the manufacturer’s closed-source library that precludes seamless conversion of raw data in vendors’ format within the web browser. With the help of increasingly performant AI tools, we expect vendors to migrate their proprietary libraries into a codebase that can be readily transpiled to multiple languages and platforms. In the meantime, without a solution to vendor proprietary libraries, the proprietary file remains the archive of record, to be reprocessed if and when needed, and Ionic serves as the fast, compact working layer, quickly converting on demand.

Several limitations need attention. The comparator values are those published by Li et al. (2018)[12] at the software versions reported therein, and some tools will have improved since then. We reran XCMS in this work for the cost comparison, but the remaining accuracy figures should be read as a benchmark of record rather than as current performance. The two tools use the CPU differently: XCMS runs one process per worker, each holding its own copy of the data, while Quant·ion runs one process with many threads that share it. On the QE HF data, XCMS could only use 4 workers before filling the 32 GB, while Quant·ion used 16 threads on the same 8-core machine. On the TripleTOF 6600, XCMS ran with 8 workers and Quant·ion with 16 threads, one per logical thread of the 8-core CPU. We could not benchmark mzMLb, its only reader relies on the HDF5 C library, which does not build for our WebAssembly target. However, highlighting this exact technical limitation is the core argument of this paper. Further, the benchmark itself consists of standard mixtures without biological matrix; recovery and false-positive counts should therefore be considered an upper bound on what could be expected from plasma, urine or tissue.

The scope of this manuscript was primarily focused on (three dimensional) LC-MS based metabolic profiling datasets, with additional proof of concept investigations conducted with ion-mobility, proteomic and imaging-MS data. To this end, any technology that uses mzML as its exchange format can benefit from this development. As such, the latter approaches and data types, as well as frontier approaches such as single-cell analysis and native mass spectrometry are the logical progression for this technology.

## Code Availability

The code underlying the software described in this study is available at: Ionic, https://github.com/phenological/ionic; Quant·ion, https://github.com/phenological/quantion; ion-beam, https://github.com/phenological/ion-beam. The command-line tool for conversion to and from the Ionic format can be installed from https://github.com/phenological/ionic/tree/main/crates/cli. Example Ionic files used by the browser application are available at https://github.com/phenological/ion-files. Code has also been deposited on Zenodo at [https://doi.org/10.5281/zenodo.22700673].

## Data Availability

The benchmark standard-mixture dataset was originally published and shared by Li et al. (2018). The proteomics dataset shown in Fig. 2 is available from PRIDE under accession PXD000001 (https://doi.org/10.6019/PXD000001), and the MALDI imaging dataset from https://doi.org/10.1186/s13742-015-0059-4. Example Ionic files used by the ion-beam browser application are available at https://github.com/phenological/ion-files. Experimental details of the targeted lipidomic and semi-targeted amino acid and biogenic amine assays are given in refs. 14,15.

## Online Methods

### 1 Raw data conversion

Raw vendor files were converted from the Bruker, SCIEX and Waters closed-source formats into the well-documented mzML format using ProteoWizard (version 3.0.25114-e35aac0). The mzML files were then converted into Ionic files (.ion), a custom binary format developed for fast data access and compact storage (see *Binary format (Ionic)* below and Supplementary Material S1). All downstream processing with Quant·ion ingests Ionic files. For comparison, the same mzML files served as input to the benchmark comparator tools as reported in the source study.

### 2 Storage footprint

#### 2.1 Internal data

The measurements reported here were made on the metabolic phenotyping archive at our centre, which at the time of writing held 103,195 converted acquisitions totalling 4.5 TB across six assays, three instrument vendors and five raw acquisition formats. Two families dominate the volume: the semi-targeted amino acid and biogenic amine assay (26,420 files, 2,412.9 GB) and the untargeted assay (21,611 files, 2,104.4 GB) together account for 99.8% of the stored bytes in 47% of the files. The remaining four assays, targeting lipids, tryptophan pathway metabolites, bile acids and short-chain fatty acids, comprise 55,164 files, more than half of the total file count, in just 11 GB, because they are multiple-reaction-monitoring acquisitions that record chromatograms rather than full spectra. Per-assay file counts and sizes are given in Supplementary Table S7.

Acquisitions originate from three vendors. Bruker instruments write a .d directory whose format is identified by the marker file it contains rather than by its name: analysis.baf for the impact II platform, and, for the timsTOF, analysis.tdf when trapped ion mobility is enabled or analysis.tsf when it is not. A census of both timsTOF instruments taken on 24 August 2026 gave 11,419 TDF against 554 TSF acquisitions: mobility-enabled acquisition is the routine mode, while TSF is largely confined to pooled QC. All acquisitions, Bruker .d, Waters .raw and SCIEX .wiff/.wiff.scan, were converted to mzML through ProteoWizard msconvert before encoding to Ionic.

These formats differ by three orders of magnitude in the number of spectra per acquisition and by roughly 500-fold in bytes per spectrum, which is what makes them a useful test of an encoding that adapts to the arrays it stores. Measured on the converted output, sampling five files per assay and nine spectra within each file, a BAF acquisition from the impact II holds 3,515 dense profile spectra of approximately 42,000 points each, at 29,558 bytes per spectrum over a 7.5 min run. A TSF acquisition from the timsTOF holds 12,636 profile spectra of approximately 8,000 points, at 20,964 bytes per spectrum over 14.0 min. A TDF acquisition from the same instrument holds 5,410,983 centroided mobility scans of 6 to 13 points, of which 44 to 66% are empty, at 54 to 60 bytes per spectrum. All carry MS1 and MS2 levels and five or six chromatograms. The impact II acquisitions mix representations, with a small number of profile spectra among otherwise centroided ones.

#### 2.2 Public data

Two public datasets were included in the format comparison to extend it beyond metabolic profiling. The first is PXD000001, the first dataset announced through ProteomeXchange and distributed by PRIDE, chosen as a data-dependent acquisition proteomics reference. In that experiment four exogenous proteins, yeast enolase, bovine serum albumin, rabbit glycogen phosphorylase and bovine cytochrome C, were spiked at defined ratios into an equimolar Erwinia carotovora lysate, digested, labelled with TMT 6-plex reagents, fractionated by reverse-phase nanoflow UPLC (nanoACQUITY, Waters) and analysed on an LTQ Orbitrap Velos (Thermo Fisher Scientific) in a Top10 HCD data-dependent method over a 60 min gradient, giving profile MS1 and centroided MS2 spectra in a single acquisition PRIDE PXD000001 ^12^. We used the deposited mzML file as distributed.

The second is the 3D MALDI imaging dataset of a mouse kidney published by Oetjen and coworkers ^13^ and deposited in MetaboLights under MTBLS176. It comprises 75 serial sections of 3.5 µm from the central part of a PAXgene-fixed, paraffin-embedded kidney, coated with sinapinic acid and acquired on a Bruker Autoflex speed in linear positive mode over 2,000 to 20,000 m/z at 50 µm lateral resolution, with 200 laser shots per spectrum. The complete dataset holds 1,362,830 profile spectra of 7,680 points each, and was exported by the original authors to imzML after user-guided rigid registration of the sections. Two properties make it useful here: 1) It is roughly two orders of magnitude larger per acquisition than any other dataset in the comparison, and 2) imzML stores one acquisition as two files, an XML metadata document and a binary sidecar linked by a shared identifier, whereas Ionic holds both in one. ProteoWizard does not support imzML, so no mzMLb equivalent could be produced, and the mzML column for this dataset is the sum of the two imzML components.

### 3 Binary Format (Ionic)

Ionic (.ion) stores the same content as mzML, spectra, chromatograms and metadata, and conversion between the two is lossless in both directions. The architecture is shown in Figure 1 and specified in full in Supplementary Material S1; the parts needed to reproduce the measurements reported here are given below. Every file begins with a fixed 1024-byte header recording the byte offset, stored length and CRC-32 checksum of every section, the number of spectra and chromatograms, the target block size and m/z window width, and the global encoding flags (endianness, compression codec, array filter). All multi-byte integers are little-endian. The file ends with the fixed 8-byte trailer \0\0\0CINOI, written after all sections have been written and the header patched, which together with the opening IONIC\0\0\0 signature confirms that the file was not truncated. Signature, trailer and all section checksums are verified before any data access.

Numeric arrays are stored as native types (16-, 32- and 64-bit integers and floats) and packed into independently compressed blocks whose uncompressed size is a configurable target recorded in the header. A block holds only segments sharing one array type, one *m/z* window and one element width, as the byte shuffle requires; the numeric type is not part of the grouping, so F64 and I64 may share a block. Two transforms are applied at different levels. Per segment, arrays of *m/z*, time or ion mobility with type F64 or F32 are delta-coded, exploiting their monotonic order, while intensity and integer arrays are stored raw. Per block, when compression is enabled and the element width is 2, 4 or 8 bytes, the block is byte-shuffled, gathering the *k*-th byte of every element into a contiguous run, then compressed with Zstandard.

Decoding reverses this order: decompression, inverse shuffle, then the inverse delta where one was applied.

Array blocks are addressed through four index sections per container. A0 is an m/z-window directory holding one entry per (window, spectrum) pair, which resolves an m/z range to the array segments that hold it. A1 is a flat array of 80-byte records in scan order, one per spectrum, holding the scalar fields used to filter spectra (retention time, base-peak m/z and intensity, selected ion m/z, total ion current, MS level, polarity, imaging coordinates), with unknown values written as NaN or zero. A2 is a flat array of 16-byte records giving the range of A3 records that each spectrum owns. A3 is a flat array of 32-byte records, each the complete address of one array segment: block identifier, starting element index and element count within the decompressed block, array type as its PSI-MS accession tail, numeric type, and the transform applied. Reading a spectrum follows the chain A1 to A2 to A3 to container, so a query resolves to a list of block identifiers before any signal is decompressed, and blocks that have no matching spectrum references are never loaded. Sections A0 - A3 are compressed on disk and decompressed in full when the file is opened; their records are fixed-width, so any record is located by arithmetic. Chromatograms use the same arrangement through sections B0 - B3, with B2 and B3 byte-identical to A2 and A3 and B1 holding a different field set of the same width.

Metadata is held in sections C, D and E, for spectra, chromatograms and global metadata respectively, as compressed columnar arrays of controlled-vocabulary parameters. Rather than encoding each parameter as a self-contained object, every property of every item is flattened into parallel columns, one column per field. Sections C and D are split into groups of up to 8192 items, each compressed as its own Zstandard frame with a directory of 32-byte entries at the section tail, so a reader that needs the parameters of one item decodes one group; section E is compressed as a single frame. Human-readable names are never stored: the CV prefix (e.g. MS, UO, NCIT) is encoded as a 1-byte enumeration code and the accession tail as a 32-bit unsigned integer, both resolved at read time against a bundled vocabulary table. Numeric values are stored as 64-bit floats and text as raw UTF-8 bytes. Section E carries a fixed 32-byte header counting items per category, each mapping one-to-one to an mzML top-level element.

## 4 Quant·ion

### 4.1 Toolkit

Peak picking is performed on one-dimensional extracted ion chromatograms (EICs), where the signal intensity is treated as a function of retention time. Peaks are detected using the Savitzky - Golay smoothing-and-derivatives approach. Smoothing is applied repeatedly using multiple window sizes, and multiple peak apexes found at similar retention times across different window sizes are treated as a single peak.

The baseline for each EIC is estimated using the adaptive iteratively reweighted penalized least squares (airPLS) method ^14^. Noise is estimated for each EIC independently. First, the EIC is scanned from start to end, then consecutive data points above the minimum value of the EIC are grouped together. For each group, the maximum intensity and the number of data points are recorded. All groups are then split into two clusters by k-means clustering. The cluster with the lowest intensity and fewest number of points is defined as noise, while the cluster with the highest intensity and highest number of points is considered to be a signal.

For each putative peak, the left and right boundaries are extended until the signal drops to the noise or baseline level, or until a valley before the next rising signal is reached. The peak area is calculated by trapezoidal integration of the signal between the final boundaries, and the peak height is defined as the maximum intensity within those boundaries. Peaks with apexes within 0.05 min of each other are merged into a single peak, and small shoulders are aggregated into the main peak area. Remaining peaks are filtered by minimum thresholds on intensity (2000 in intensity), peak width (5 points), and signal-to-noise ratio (3). These threshold parameters were fixed across all datasets.

### 4.2 External benchmark dataset

For this study, we used the benchmark standard-mixture dataset published by Li and coworkers ^15^, which provides a defined chemical ground truth for evaluating untargeted metabolic profiling feature detection algorithms. Two mixtures, SA and SB, were prepared from 1,100 drugs and metabolites of 100-1300 Da and diluted in acetonitrile/water (1:1, v/v); the samples contain no biological matrix. Of these, 970 compounds were present at equal concentration in both mixtures and form a constant, non-differential background (the “matrix group” of Li et al. (2018)), while the remaining 130 were spiked at defined SB:SA ratios (1:16, 1:4, 1:2, 2:1, 4:1 and 16:1) across six differential groups, constituting a set of known discriminating markers. Both mixtures were acquired in positive ion mode on two high-resolution platforms, an SCIEX TripleTOF 6600 and a Thermo Q Exactive HF Orbitrap, in four replicates and five replicates, respectively. Since not every compound ionises and is detected, and because the two platforms have different resolution, targeted extraction confirmed a single feature for only a subset of the 1100 inputs: 970 features on the TripleTOF (867 background, 103 differential) and 836 on the QE HF (760 background, 76 differential). Applying the source paper’s marker criteria (fold change >2 or <0.5, p < 0.05) to the differential features yields benchmark discriminating-marker lists of 68 and 50 features, respectively. Feature recovery and marker selection in this work are scored against these confirmed, platform-specific lists rather than against the 1100 input compounds. Note that the TripleTOF benchmark feature count coincides numerically with the number of non-differential compounds; the two sets are distinct.

We ran Quant·ion on this benchmark. The values reported for the comparator tools, MarkerView (SCIEX, Framingham, MA, USA), MS-DIAL ^8^, MZmine 2 ^11^, XCMS ^10^, and Compound Discoverer (Thermo Fisher Scientific, Waltham, MA, USA), are taken directly from ref^15^, at the versions reported therein; Quant·ion and XCMS were additionally run in this work.

### 4.3 Sample-level feature detection

Untargeted feature detection was performed independently on each sample, across the full *m/z* and retention-time space. A uniform *m/z* grid was built from *m/z* 100 to 1300 with a step size of 0.005 Da. For each grid point, an EIC was extracted spanning 0.4 to 36.0 minutes, using a tolerance of 10 ppm or 0.003 Da, whichever is larger; the absolute floor prevents the ppm window from becoming impractically narrow at low *m/z*. Peak picking was then applied to each EIC as described above. For each putative peak, the *m/z* value was refined by selecting the centroid of highest intensity within the local *m/z* window, and duplicate candidate masses were removed. Final EICs were recalculated for each refined *m/z* using a wider tolerance of 20 ppm or 0.005 Da whichever is larger, accommodating the residual mass dispersion of the pooled candidates, and re-picked to produce the definitive feature list. Each feature is identified by its *m/z* and carries the following attributes: retention time, median intensity, and median peak area, summarised across the samples in which it was detected.

### 4.4 Benchmark scoring

Each entry in the benchmark feature lists of Li et al. (2018) (Tables S7 and S8 in the Supporting Information of Li et al. (2018); 970 and 836 entries) was scored as found when a consensus feature fell within the limits used in that study: mass error below 10 ppm and retention-time shift below 0.3 min. Some benchmark entries share the same *m/z* and retention time, so they match the same feature and all of them are counted, since the pipelines do not report duplicates. The TripleTOF 6600 list contains 44 such positions, 43 holding two compounds and one holding three, 89 compounds in total, whose masses differ by 0.0006 Da on average (maximum 0.0016 Da), far below the resolution of the instrument; at those positions Quant·ion reported a single feature at 43 of the 44 positions, including the one holding three compounds, and XCMS at 41 of 44. The QE HF list has no such pairs. The same rule was applied to the XCMS results in Figure 5a.

All measurements were made on a 2019 Mac Pro (3.5 GHz 8-core Intel Xeon W, 32 GB RAM), on CPU only. Quant·ion ran through its R wrapper (v0.1.0) on Ionic files, using 10 threads on both datasets. XCMS (v4.8.0) ran on the same acquisitions, converted to mzML and centroided first, using 8 workers on the TripleTOF 6600 and 4 on the QE HF. XCMS workers run in parallel, and each one keeps its own copy of a sample in memory. QE HF samples hold more MS1 spectra, and each spectrum holds more points, so running 8 at once will overload the 32 GB RAM. The XCMS times include centroiding needed to use CentWave, Ionic files do not need this step. We recorded the time each call took, and the most memory R and its worker processes held at once, sampled once per second. Each value comes from a single run.

### 4.5 Biological-level feature detection

Features from all samples were combined and grouped across the full dataset. Features within 10 ppm or 0.0025 Da, whichever is larger, and 0.3 min of each other were assigned to the same cluster. In cases where the same sample contributed more than one feature to a cluster, only the feature with the highest intensity was considered. Gap filling was performed for any sample with no feature in a given cluster, i.e., the raw data was searched again using the cluster *m/z* and retention-time window to recover signals missed in the initial detection iteration. For each cluster, the final reported *m/z*, retention time, intensity, and area were calculated as the median values across all samples in which a feature was detected. The number of samples in which each feature was detected was also recorded. Peak areas were log2-transformed and compared between SB and SA with Welch’s t-test. A feature was counted as a marker when its fold change was above 2.0 or below 0.5, and P was below 0.05.

### 4.6 Implementation

The Ionic file-format library and the Quant·ion toolkit were written in Rust and released as two separate open-source projects under the MIT license. Ionic is a Rust library that handles parsing of mzML files, encoding and decoding of the Ionic format, and management of the resultant data. It can be used as a library or driven through a command-line interface (CLI). Quant·ion is the data-processing toolkit built on top of Ionic, delivering the functions evaluated in this work: peak picking, baseline correction, noise estimation, and untargeted feature detection.

Quant·ion exposes its processing functions to Python, R, and JavaScript/TypeScript users through dedicated language wrappers, so that the same underlying Rust source is used across environments, and the results do not depend on the language. Ionic and the mzML parser are used as dependencies of Quant·ion. The Python and R wrappers can be installed directly from the GitHub repository and the JavaScript wrapper from npm; pre-built binaries are provided for Linux and macOS on x86-64 and arm64, and for Windows on x86-64. No external system libraries (such as HDF5) are required at runtime; the Zstandard codec is statically linked into the pre-built binaries.

Peak detection relies on Savitzky - Golay smoothing and its derivatives, and is built on three components that were reimplemented directly in Rust from the corresponding mljs libraries: smoothing uses ml-savitzky-golay-generalized ^16^, baseline correction uses airPLS from ml-airpls ^17^, and peak fitting uses non-linear (Levenberg - Marquardt) curve fitting from ml-levenberg-marquardt ^18^. These are build-time implementations providing the algorithms, not runtime dependencies of the format. Internal helpers handle the surrounding processing logic.

## Acknowledgements

JW acknowledges support from Universidad del Valle. All authors thanks the invaluable support of Bruker BBIO and BDAL.

## Contributors

JOM and JW designed the Ionic format. JOM and JOM implemented the Rust core; JOM the R, Python and JavaScript wrappers, and the ion-beam web application. JOM and JW performed the benchmark evaluation. NGL, MC, SS and LW acquired and curated the internal datasets. JKN. and EH secured funding and contributed to interpretation. JW and JOM wrote the manuscript with input from all authors. All authors read and approved the final version.

## Supplementary Information

The online version contains supplementary material available at [DOI]. Supplementary Information (PDF): Supplementary Note 1, the Ionic file format specification, with Supplementary Tables 1–5 and Supplementary Fig. 1; Supplementary Tables 6 and 7, query cost across formats and composition of the phenotyping archive; Supplementary Note 2, the 44 isobaric co-eluting positions in the TripleTOF 6600 benchmark list, with Supplementary Table 8.

