## Supplementary Information for "A dependency-free, streamable format and cross-language toolkit for scalable LC-MS feature detection: reading only what you need"

**Supplementary material	1**

S1- Ionic	3

File Header	3

Spectrum sections (A0, A1, A2, A3)	5

Section C, D and E	7

File Trailer	10

S2 - Isobaric + Co-eluting compounds	11

### S1- Ionic

Every Ionic file begins with a fixed 1024-byte header. Because the header is fixed-size, a parser can read it with a single system call and immediately compute the byte position of any section without scanning the file. The header stores: the byte offset and byte length of every section, the total number of spectra and chromatograms, per-section CRC-32 integrity checksums, and global encoding flags (endianness, compression codec, array filter) (**Table S1**). All multi-byte integers are stored in little-endian byte order.

#### File Header

**Table S1. File header field layout.**

| Offset (B) | Size (B) | Field | Type | Description |
| --- | --- | --- | --- | --- |
| 0 | 8 | file_signature | u8[8] | IONIC\0\0\0. |
| 8 | 1 | endianness_flag | u8 | 0 = little-endian. Must be 0; every integer in the file is little-endian. |
| 9 | 2 | format_version | u16 | Format version. |
| 11 | 1 | compression_codec | u8 | Compression codec: 0 = none, 1 = zstd. |
| 12 | 1 | compression_level | u8 | Zstandard compression level (0-22). |
| 13 | 1 | default_array_filter | u8 | Transform applied at the block level: 0 = raw, 1 = byte shuffle, 2 = delta. |
| 14 | 2 | reserved | u8[2] | Reserved. |
| 16 | 8 | target_block_uncompressed_bytes | u64 | Target uncompressed size of one array block. |
| 24 | 4 | target_mz_window | u32 | m/z window width in m/z units. |
| 28 | 4 | reserved | u8[4] | Reserved. |
| 32 | 8 | off_spec_window_directory (A0) | u64 | Offset of the spectrum m/z-window directory. |
| 40 | 8 | len_spec_window_directory (A0) | u64 | Stored length, compressed when the codec is zstd. |
| 48 | 8 | off_spec_summary (A1) | u64 | Offset of the spectrum fast filter summary. |
| 56 | 8 | len_spec_summary (A1) | u64 | Stored length. |
| 64 | 8 | off_spec_entries (A2) | u64 | Offset of the spectrum array index. |
| 72 | 8 | len_spec_entries (A2) | u64 | Stored length. |
| 80 | 8 | off_spec_array_addresses (A3) | u64 | Offset of the spectrum array address table. |
| 88 | 8 | len_spec_array_addresses (A3) | u64 | Stored length. |
| 96 | 8 | off_chrom_window_directory (B0) | u64 | Offset of the chromatogram window directory. |
| 104 | 8 | len_chrom_window_directory (B0) | u64 | Stored length. |
| 112 | 8 | off_chrom_summary (B1) | u64 | Offset of the chromatogram summary. |
| 120 | 8 | len_chrom_summary (B1) | u64 | Stored length. |
| 128 | 8 | off_chrom_entries (B2) | u64 | Offset of the chromatogram array index. |
| 136 | 8 | len_chrom_entries (B2) | u64 | Stored length. |
| 144 | 8 | off_chrom_array_addresses (B3) | u64 | Offset of the chromatogram array address table. |
| 152 | 8 | len_chrom_array_addresses (B3) | u64 | Stored length. |
| 160 | 8 | off_spec_meta (C) | u64 | Offset of the spectrum metadata section. |
| 168 | 8 | len_spec_meta (C) | u64 | Length of the section. |
| 176 | 8 | off_chrom_meta (D) | u64 | Offset of the chromatogram metadata section. |
| 184 | 8 | len_chrom_meta (D) | u64 | Length of the section. |
| 192 | 8 | off_global_meta (E) | u64 | Offset of the global metadata section. |
| 200 | 8 | len_global_meta (E) | u64 | Length of the section. |
| 208 | 8 | off_spec_container | u64 | Offset of the spectrum array-block container. |
| 216 | 8 | len_spec_container | u64 | Length of the spectrum container. |
| 224 | 8 | off_chrom_container | u64 | Offset of the chromatogram array-block container. |
| 232 | 8 | len_chrom_container | u64 | Length of the chromatogram container. |
| 240 | 8 | spec_block_count | u64 | Number of blocks in the spectrum container. |
| 248 | 8 | chrom_block_count | u64 | Number of blocks in the chromatogram container. |
| 256 | 8 | spectrum_count | u64 | Number of spectra; also the A1 and A2 record counts. |
| 264 | 8 | chrom_count | u64 | Number of chromatograms; also the B1 and B2 record counts. |
| 272 | 8 | spec_array_type_count | u64 | Distinct array types across spectra. |
| 280 | 8 | chrom_array_type_count | u64 | Distinct array types across chromatograms. |
| 288 | 8 | spec_meta_count | u64 | Total metadata rows in C, the sum over all its groups. |
| 296 | 8 | spec_meta_numeric_count | u64 | Total numeric values in C. |
| 304 | 8 | spec_meta_string_count | u64 | Total string values in C. |
| 312 | 8 | chrom_meta_count | u64 | Total metadata rows in D. |
| 320 | 8 | chrom_meta_numeric_count | u64 | Total numeric values in D. |
| 328 | 8 | chrom_meta_string_count | u64 | Total string values in D. |
| 336 | 8 | global_meta_count | u64 | Total metadata rows in E. |
| 344 | 8 | global_meta_numeric_count | u64 | Total numeric values in E. |
| 352 | 8 | global_meta_string_count | u64 | Total string values in E. |
| 360 | 8 | spec_meta_uncompressed_bytes | u64 | Decompressed size of Section C. |
| 368 | 8 | chrom_meta_uncompressed_bytes | u64 | Decompressed size of Section D. |
| 376 | 8 | global_meta_uncompressed_bytes | u64 | Decompressed size of Section E. |
| 384 | 8 | plain_len_spec_window_directory (A0) | u64 | Uncompressed length of A0. |
| 392 | 8 | plain_len_chrom_window_directory (B0) | u64 | Uncompressed length of B0. |
| 400 | 8 | total_file_size | u64 | Total file size in bytes. The reader locates the trailer at total_file_size - 8. |
| 408 | 8 | meta_group_size | u64 | Items per metadata group in C and D (8192). |
| 416 | 8 | spec_meta_group_count | u64 | Number of groups in Section C. |
| 424 | 8 | chrom_meta_group_count | u64 | Number of groups in Section D. |
| 432 | 536 | reserved | u8[536] | Reserved. |
| 968 | 4 | spec_window_directory_crc32 (A0) | u32 | CRC-32 of Section A0 as stored. |
| 972 | 4 | spec_summary_crc32 (A1) | u32 | CRC-32 of Section A1 as stored. |
| 976 | 4 | spec_entries_crc32 (A2) | u32 | CRC-32 of Section A2 as stored. |
| 980 | 4 | spec_array_addresses_crc32 (A3) | u32 | CRC-32 of Section A3 as stored. |
| 984 | 4 | chrom_window_directory_crc32 (B0) | u32 | CRC-32 of Section B0 as stored. |
| 988 | 4 | chrom_summary_crc32 (B1) | u32 | CRC-32 of Section B1 as stored. |
| 992 | 4 | chrom_entries_crc32 (B2) | u32 | CRC-32 of Section B2 as stored. |
| 996 | 4 | chrom_array_addresses_crc32 (B3) | u32 | CRC-32 of Section B3 as stored. |
| 1000 | 4 | spec_directory_crc32 | u32 | CRC-32 of the spectrum block directory. |
| 1004 | 4 | chrom_directory_crc32 | u32 | CRC-32 of the chromatogram block directory. |
| 1008 | 4 | spec_meta_crc32 (C) | u32 | CRC-32 of Section C. |
| 1012 | 4 | chrom_meta_crc32 (D) | u32 | CRC-32 of Section D. |
| 1016 | 4 | global_meta_crc32 (E) | u32 | CRC-32 of Section E. |
| 1020 | 4 | header_crc32 | u32 | CRC-32 over header bytes 0-1019. |

##### Spectrum sections (A0, A1, A2, A3)

Spectral data is organised across four index sections and one compressed data container. Reading a spectrum follows a strict four-step chain: the parser 1) scans Section A1 to identify which spectra match the query; 2) reads the corresponding Section A2 record to find the spectrum's slice of Section A3; 3) reads those Section A3 records to obtain the exact block, element offset, and element count for each array segment and 4) decompresses only the blocks it needs from the spectrum container. Every block that is not referenced by the query remains compressed and is never loaded. Sections A1, A2 and A3 are compressed on disk and decompressed in full when the file is opened. Their records are fixed-width, so the position of any record is a single arithmetic calculation. This workflow is illustrated with a worked example in Figure 3.

**Section A0: m/z-window directory (variable length).** A0 resolves an *m/z* range to the array segments that hold it (Figure 3). The *m/z* axis is divided into fixed-width windows, the window of an *m/z* value being floor(*m/z* $\div$ w), where w is the integer width stored in the header at offset 24. A0 holds one entry per (window, spectrum) pair, for every spectrum with *m/z* values in that window.

The section consists of two counts followed by four u32 arrays, one after another (**Table S2**): starts has window_count + 1 values; the other three have entry_count values each and are indexed together. Window b owns entries from starts[b] to starts[b+1] − 1. Each entry gives the index of the spectrum it belongs to (spectrum_index) and two A3 record indices, one for the *m/z* segment (mz_address) and one for the intensity segment (intensity_address). Those records hold the block, element offset, element count, and dtype; A0 stores none of these. Retention time comes from A1[spectrum_index]. A query over [lo, hi] visits windows from floor(lo / w) to floor(hi / w), and decodes only the blocks their entries reference. Within a window, spectrum_index ascends so that a single spectrum can be found by binary search.

**Table S2. Section A0 field layout.**

| Offset (B) | Size (B) | Field | Type | Description |
| --- | --- | --- | --- | --- |
| 0 | 4 | window_count | u32 | Number of windows. |
| 4 | 4 | entry_count | u32 | Number of entries. |
| 8 | 4(w_c + 1) | starts | u32[w_c + 1] | starts[b]..starts[b+1] is window b's entry slice. |
| 12 + 4w_c | 4e_c | spectrum_index | u32[e_c] | Owning spectrum. |
| 12 + 4(w_c + e_c) | 4e_c | mz_address | u32[e_c] | A3 record of the m/z segment. |
| 12 + 4(w_c + 2e_c) | 4e_c | intensity_address | u32[e_c] | A3 record of the intensity segment. |

w_c = window_count, e_c = entry_count. Offsets are within the decompressed section; total length is 12 + 4(w_c + 3e_c) bytes

**Section A1: Spectra fast filter summary (80 bytes per spectrum).** A1 lets a query discard spectra before any array data is read. It is a flat array of 80-byte records in scan order, one per spectrum, so the record for spectrum i begins at off_spec_summary + i × 80. It holds the scalar fields most commonly used to filter spectra, all as native numeric types (**Table S3**). Unknown values are IEEE 754 NaN in the floating-point fields and zero in the integer fields. Finite rt values ascend from record to record, so when every spectrum has an rt, a retention-time range can be found by binary search.

**Table S3. Section A1 field layout.**

| Offset (B) | Size (B) | Field | Type | Description |
| --- | --- | --- | --- | --- |
| 0 | 8 | rt | f64 | Retention time at scan start (PSI-MS: MS:1000016), in the unit given by rt_unit at offset 54. No unit normalisation is performed. NaN = unknown. |
| 8 | 8 | base_peak_mz | f64 | m/z of the most intense peak (MS:1000504). NaN = unknown. |
| 16 | 8 | selected_ion_mz | f64 | Precursor ion m/z (MS:1000744); populated for MS² and above. NaN = unknown. |
| 24 | 8 | base_peak_int | f64 | Intensity of the most intense peak (MS:1000505). NaN = unknown. |
| 32 | 8 | total_ion_current | f64 | Sum of all ion intensities in the scan (MS:1000285). NaN = unknown. |
| 40 | 1 | ms_level | u8 | MS level (MS:1000511). 0 = unknown. |
| 41 | 1 | polarity | u8 | Acquisition polarity: 0 = unknown, 1 = positive (MS:1000130), 2 = negative (MS:1000129). |
| 42 | 4 | x | u32 | Scan position X for imaging data. 0 = unknown. |
| 46 | 4 | y | u32 | Scan position Y for imaging data. 0 = unknown. |
| 50 | 4 | z | u32 | Scan position Z for imaging data. 0 = unknown. |
| 54 | 1 | rt_unit | u8 | Unit of the rt field: 0 = other, 1 = second, 2 = minute, 3 = millisecond. |
| 55 | 25 | reserved | u8[25] | Reserved; must be zero. Accommodates future fields without changing the record size. |

**Section A2: Spectrum array index (16 bytes per spectrum).** A2 maps a spectrum to its array segments. It is a flat array of 16-byte records, one per spectrum in scan order (**Table S4**), so the record for spectrum i begins at off_spec_entries + i × 16. Each record holds two numbers: where the first record of the spectrum starts in A3 (arr_ref_start) and how many records it owns (arr_ref_count). Spectrum i owns A3 records from arr_ref_start to arr_ref_start + arr_ref_count − 1. A spectrum has one segment per array per *m/z* window, and the format assumes no fixed count. A2 stores nothing else: the block, element offset, element count and dtype come from those A3 records.

**Table S4. Section A2 field layout.**

| Offset (B) | Size (B) | Field | Type | Description |
| --- | --- | --- | --- | --- |
| 0 | 8 | arr_ref_start | u64 | Index of the first row in Section A3 that belongs to this spectrum. |
| 8 | 8 | arr_ref_count | u64 | Number of consecutive Section A3 records belonging to this spectrum. |

**Section A3: Array address table (32 bytes per array segment).** A3 is a flat array of 32-byte records (Table S5). Each record is the complete address of one array segment: the starting element index within the decompressed block (off_element), the number of elements (len_element), the block it lives in (block_id), the PSI-MS accession tail identifying the array type (array_type, e.g. 1000514 for m/z, 1000515 for intensity), the numeric type of the elements (dtype: 1 = F64, 2 = F32, 3 = F16, 4 = I16, 5 = I32, 6 = I64), and the pre-compression transform applied to the segment (array_filter: 0 = raw, 1 = byte shuffle, 2 = delta). Three further fields complete the record: encoded_len (0 for fixed-width arrays, otherwise the stored byte count), continues_previous_segment (1 when the segment continues the array in the record above, 0 when it starts a new one), and array_cv_code (the controlled vocabulary of array_type; 0 = MS). In Figure 3, spectrum 102 owns eight records, A3[816] to A3[823], because its *m/z* and intensity arrays are each split across four *m/z* windows. For the 0 - 250 Da window, these are A3[816] (m/z, F64, block 0, element offset 150, 200 elements) and A3[820] (intensity, F32, block 4, element offset 150, 200 elements).

**Table S5. Section A3 field layout.**

| Offset (B) | Size (B) | Field | Type | Description |
| --- | --- | --- | --- | --- |
| 0 | 8 | off_element | u64 | Starting element index inside the decompressed block. In elements, not bytes. |
| 8 | 8 | len_element | u64 | Number of elements in this segment. |
| 16 | 4 | block_id | u32 | Block in the container that holds the segment. |
| 20 | 4 | array_type | u32 | PSI-MS accession tail of the array type; 1000514 = m/z, 1000515 = intensity. |
| 24 | 1 | dtype | u8 | Numeric type of the elements: 1 = F64, 2 = F32, 3 = F16, 4 = I16, 5 = I32, 6 = I64. |
| 25 | 1 | array_filter | u8 | Transform applied to this segment: 0 = raw, 1 = byte shuffle, 2 = delta. |
| 26 | 4 | encoded_len | u32 | 0 for fixed-width arrays; otherwise the stored byte count of the segment. |
| 30 | 1 | continues_previous_segment | u8 | 0 when the segment starts a new array, 1 when it continues one already started. |
| 31 | 1 | array_cv_code | u8 | Controlled vocabulary of array_type; 0 = MS |

The spectrum container stores all numeric arrays as independently compressed blocks. Each block holds only segments sharing one array type, one *m/z* window, and one element width (1, 2, 4 or 8 bytes), as the byte-shuffle requires. Dtype is not part of the grouping; F64 and I64 may share a block. Segments from many spectra share a block; what varies per spectrum is the element slice given by off_element and len_element. Two transforms are applied at different levels. Per segment, arrays of type *m/z*, time or ion mobility with dtype F64 or F32 are delta-coded, exploiting their monotonic order; intensity and integer arrays are stored raw. Per block, when compression is enabled and the element width is 2, 4 or 8 bytes, the block is byte-shuffled, the k-th byte of every element gathered into a contiguous run, then Zstandard-compressed. Decoding reverses the order: Zstandard decompression, inverse byte-shuffle, then the inverse delta where one was applied. Blocks that are not referenced by any queried spectrum remain compressed and are never loaded; in **Figure 3**, blocks 0, 1, 4, and 5 are decoded, the *m/z* and intensity blocks of the two windows overlapping the query, while blocks 2, 3, 6, and 7 remain compressed.

The same four-section arrangement rules chromatogram data through Sections B0 (window directory), B1 (summary), B2 (array index), B3 (array address table), and the chromatogram container. B2 and B3 are byte-identical to A2 and A3. B1 is also 80 bytes per item but holds a different field set: lowest_mz (f64, offset 0), highest_mz (f64, 8), lowest_wavelength (f64, 16), highest_wavelength (f64, 24), lowest_ion_mobility (f64, 32), highest_ion_mobility (f64, 40) and polarity (u8, 48), with the remainder reserved; NaN = unknown. Chromatograms are not split by window; B0 holds a single window 0 with one entry per chromatogram. B0 reuses A0's column layout, but its mz_address column points to the chromatogram's time array rather than an *m/z* array.

##### Section C, D and E

Sections C, D, and E store spectrum metadata, chromatogram metadata, and global metadata respectively. All three use the same structure. Each CV parameter is stored as one row across twelve parallel arrays: a Contiguous Index (CI) that gives the row range for each item; tag, owner, and parent identifiers (MTI, MOI, MPI); ontology reference (MRI, e.g. 0 = MS, 1 = UO); CV accession tail (MAN, e.g. 1000514 for MS:1000514); unit reference and accession (MURI, MUAN); value kind (VK: 0 = number, 1 = string, 2 = absent); a numeric pool (VN, 64-bit floats); a string length array (VLEN); and a UTF-8 string pool (VS). Two further columns are not stored but computed on read: the value index (VI) and the string offset array (VOFF). Values go into their pools in row order and are never reused, so the first numeric row takes VN[0], the second VN[1], and so on; VI is simply how many values of that kind have been seen so far. VOFF is the running total of VLEN, giving the byte at which each string starts in VS. To read the metadata for item *i*: read CI[*i*] and CI[*i*+1] to get the row range, then for each row use VK to pick the value from VN or VS. Section E also has a 32-byte general header that counts items per category, each mapping 1-to-1 to an mzML top-level element (*<fileDescription>*, *<sampleList>*, *<instrumentConfigurationList>*, etc.). Attributes with no PSI-MS equivalent use the ATTR namespace.

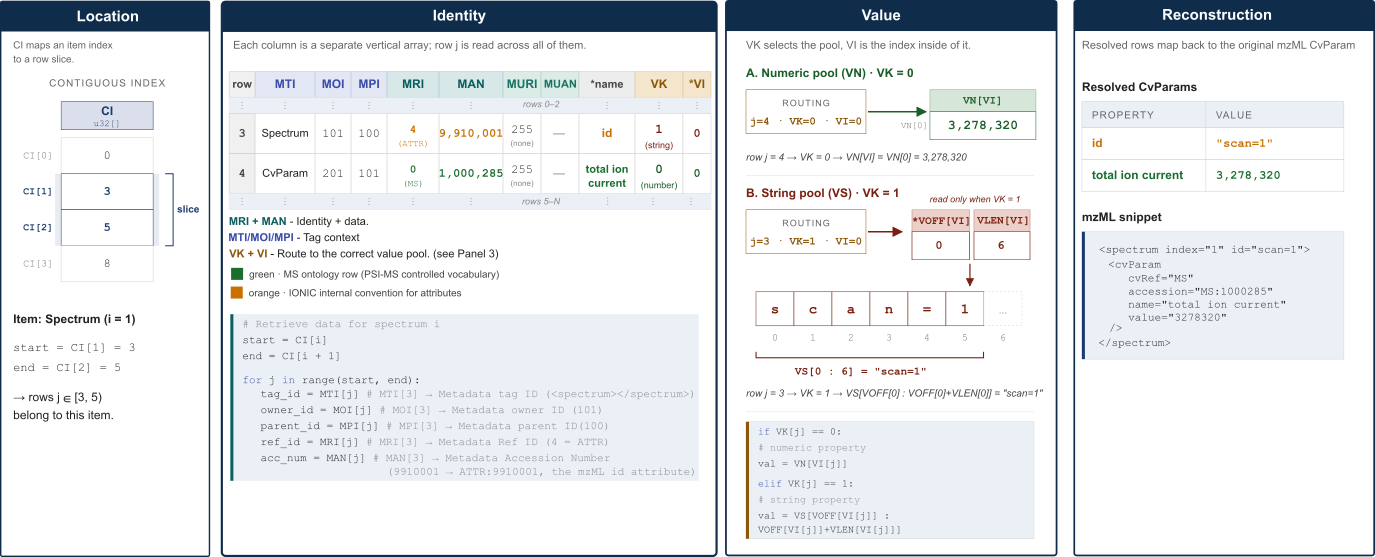

**Figure S2.** Schema for columnar metadata storage and decoding in Sections C and D of the Ionic format. The Contiguous Index (CI) maps each spectrum or chromatogram to a row slice in a set of parallel column arrays, where each row encodes one CV parameter through its identity (MRI, MAN), tag context (MTI, MOI, MPI), and value kind (VK). Numeric values are read directly from the floating-point pool (VN), while string values are resolved through offset and length indices into a concatenated byte pool (VS). The rightmost panel demonstrates full round-trip reconstruction from raw column data back to a valid mzML CvParam. Section C stores spectrum metadata; Section D is identical in structure and stores chromatogram metadata. *Columns marked * are not stored in the file but reconstructed when the file is read: name from MRI + MAN against the controlled vocabulary, and VI as a per-kind running ordinal over VK. The string offset array VOFF, shown in the Value panel, is derived as well: the running total of VLEN.*

Sections C, D, and E store metadata as compressed columnar arrays. Instead of encoding each CV parameter as a self-contained object, the format flattens every property of every item into a set of parallel columns where each column holds one field across all rows. Sections C and D are split into groups of up to 8192 items, each compressed as its own Zstandard frame, with a directory of 32-byte entries at the section tail; each group also begins with a 12-byte header giving its row, numeric and string counts. Section E is not split and is compressed as one frame. A reader that wants one item's metadata decodes only the group holding it. The complete on-disk column order is fixed:

CI | MOI | MPI | MTI | MRI | MAN | MURI | MUAN | VK | VN | VLEN | VS

In Sections C and D, the Contiguous Index (CI) is per group: a u32 array with one entry per item in the group plus a final entry, and row numbers counted from the group's first row. It works as a range map: item i of the group owns all rows between CI[i] (inclusive) and CI[i+1] (exclusive). In **Figure S2**, spectrum i = 1 has CI[1] = 3 and CI[2] = 5, so rows 3 and 4 belong to that spectrum. The final entry equals that group's meta_count, given in its 12-byte header. Section E is not split, so its CI spans the whole section: item_count + 1 entries, the last equal to the section's total number of rows.

Each row encodes one CV parameter or XML attribute. The columns that describe what the row is are as follows: MTI (u8 tag context, e.g. Spectrum, CvParam), MOI (owner ID, u32), MPI (parent ID, u32), MRI (Cv ref code: 0 = MS, 1 = UO, 2 = NCIT, 3 = PEFF, 4 = ATTR, 5 = IMS, 255 = none, u8), and MAN (numeric accession tail, e.g. 1000285 for MS:1000285, u32). Unit information follows the same pattern through MURI and MUAN. In Sections C and D the ids are item-local: the counter restarts at 2 for each item, and id 1 is reserved for the shared list node. A reader must rebase them before the parent - child links are usable, compute stride = 1 + max(MOI, MPI) over the group, then map every id other than 1 to item_index × stride + local_id. Section E stores global ids and needs no rebasing.

The value of each row is given by two fields: VK (u8, Value Kind) selects the pool, and VI (u32, Value Index) is the index inside it. VK = 0 means numeric; the value is VN[VI[j]], a 64-bit float. VK = 1 means string; the value is the byte slice VS[VOFF[VI[j]]: VOFF[VI[j]] + VLEN[VI[j]]]. VK = 2 means the property has no value. In **Figure S2**, row 3 has VK = 1, VI = 0, routing to VS[0:6] = "scan=1"; row 4 has VK = 0, VI = 0, routing to VN[0] = 3,278,320. Both together reconstruct the id attribute and the total ion current CV parameter of that spectrum.

Human-readable names are never stored in the file. The accession number in MAN is resolved at read time against a standard CV table: MAN = 1000285 with MRI = 0 yields MS:1000285, looked up as "total ion current".

Section C holds spectrum metadata; the header's spectrum_count and metadata row count are the file-wide totals, equal to the sums over C's groups. Section D holds chromatogram metadata with the analogous header counts. Both sections are otherwise structurally identical.

Section E holds global metadata file description, sample, instrument configuration, software, data processing, and run information. It begins with a fixed 32-byte header before the column data, holding nine u16 counts (one per mzML top-level category, in the order: fileDescription, referenceableParamGroupList, sampleList, instrumentConfigurationList, softwareList, dataProcessingList, scanSettingsList, cvList, run) followed by 14 reserved bytes. The CI that follows indexes through the concatenation of all categories in that fixed order, so item_count for Section E is the sum of all nine counts.

#### File Trailer

The last 8 bytes of every valid Ionic file are the fixed ASCII sequence \0\0\0CINOI. Together with the opening IONIC\0\0\0 signature, this trailer allows a parser to confirm before reading any section that the file was written completely and was not truncated while writing. The encoder must write the trailer as the final operation, after all sections have been written and after the header has been patched with all offsets, counts, and checksums.

**Table S6.** Cost of opening one acquisition and extracting a single extracted-ion chromatogram, and the resulting signal, for the same acquisitions stored as mzML, mzMLb and Ionic. Each row is one format read by the implementation available for it; no single implementation reads all three, so the timings reflect reader and format together. Times in milliseconds.

| **Dataset** | **Target compound** | ***m/z*** | **RT (min)** | **Format / reader** | **Open (ms)** | **EIC (ms)** | **Total (ms)** | **Max intensity** |
| --- | --- | --- | --- | --- | --- | --- | --- | --- |
| iron_is | L-Phenylalanine-¹³C₉,¹⁵N | 176.1135 | 3.97 | Ionic / Quant·ion | 9.7 | 5.1 | **14.8** | 44154 |
|  |  |  |  | mzML / pyOpenMS | 497 | 13.6 | **511** | 44154 |
|  |  |  |  | mzMLb / pyteomics + pyOpenMS | 3037 | 12.9 | **3050** | 44154 |
| aa_is | Lysine-¹³C₆,¹⁵N₂ | 248.1157 | 4.22 | Ionic / Quant·ion | 7.7 | 3.7 | **11.5** | 10098 |
|  |  |  |  | mzML / pyOpenMS | 508 | 41.9 | **550** | 10098 |
|  |  |  |  | mzMLb / pyteomics + pyOpenMS | 2108 | 25.7 | **2134** | 10098 |
| MTBLS733 | T0226 | 192.1385 | 13.76 | Ionic / Quant·ion | 20.9 | 14.5 | **35.4** | 5.32792770 × 10⁸ |
|  |  |  |  | mzML / pyOpenMS | 2831 | 337 | **3168** | 5.32792770 × 10⁸ |
|  |  |  |  | mzMLb / pyteomics + pyOpenMS | 9785 | 496 | **10281** | 5.32792770 × 10⁸ |

Open is the time required before the first query can be answered; EIC is the extraction that follows. Max intensity is the maximum of the extracted chromatogram. The first two compounds are stable-isotope-labelled internal standards of known m/z and retention time, so agreement of their maxima across formats confirms identity rather than coincidence. In the aa_is assay, amino acids are derivatized with AccQ-Tag; lysine carries two labels and is measured as [M+2H]2+. Chromatograms were extracted at 20 ppm or 0.005 Da, whichever is larger, on the workstation given in Online Methods, and each value is a single measurement. Ionic files were read with Quant·ion 0.1.0 through its Python wrapper. mzML files were read and processed with pyOpenMS 3.5.0. mzMLb files required pyteomics 5.0.1 to read the file and pyOpenMS 3.5.0 to compute the chromatogram. mzMLb stores native numeric types and would be expected to open faster than mzML; these timings reflect the composite reader rather than the format.

**Table S7.** Composition of the in-house metabolic phenotyping archive at the time of writing. Sizes are of the converted Ionic files.

| **Assay** | **Raw format(s)** | **Files** | **Total (GB)** | **Mean (MB)** | **Median (MB)** | **Max (MB)** |
| --- | --- | --- | --- | --- | --- | --- |
| Semi-targeted amino acids and biogenic amines | BAF | 26,420 | 2,412.9 | 93.5 | 90.4 | 407.3 |
| Untargeted | BAF, TDF, TSF | 21,611 | 2,104.4 | 99.7 | 68.9 | 814.0 |
| Targeted lipids | WIFF | 39,940 | 9.1 | 0.2 | 0.2 | 5.5 |
| Targeted tryptophan pathway | RAW | 11,964 | 1.6 | 0.1 | 0.1 | 0.2 |
| Targeted bile acids | RAW | 2,181 | 0.3 | 0.1 | 0.1 | 0.3 |
| Targeted short-chain fatty acids | RAW | 1,079 | <0.1 | <0.1 | <0.1 | <0.1 |
| **Total** |  | **103,195** | **4,528.3** |  |  |  |

The two untargeted timsTOF formats are distinguished by the marker file inside the vendor .d directory: analysis.tdf with trapped ion mobility enabled, analysis.tsf without. A census on 24 August 2026 gave 11,419 TDF and 554 TSF acquisitions. WIFF denotes SCIEX and RAW denotes Waters acquisitions, both converted through ProteoWizard; Bruker acquisitions were read through the vendor SDKs and written directly to Ionic.

### S2 - Isobaric + Co-eluting compounds

| mz | rt | ids | n |
| --- | --- | --- | --- |
| 442.2233 | 3.24 | TTOF0778 (m/z: 442.2233), TTOF0942 (m/z: 442.2234) | 2 |
| 313.1767 | 3.39 | TTOF0229 (m/z: 313.1770), TTOF0767 (m/z: 313.1765) | 2 |
| 321.0981 | 3.77 | TTOF0373 (m/z: 321.0981), TTOF0721 (m/z: 321.0980) | 2 |
| 337.2124 | 3.78 | TTOF0034 (m/z: 337.2121), TTOF0784 (m/z: 337.2127) | 2 |
| 499.2264 | 4.06 | TTOF0522 (m/z: 499.2260), TTOF0711 (m/z: 499.2268) | 2 |
| 400.1474 | 4.07 | TTOF0062 (m/z: 400.1475), TTOF0865 (m/z: 400.1473) | 2 |
| 363.1454 | 4.27 | TTOF0402 (m/z: 363.1452), TTOF0448 (m/z: 363.1456) | 2 |
| 400.1472 | 4.68 | TTOF0776 (m/z: 400.1473), TTOF0866 (m/z: 400.1472) | 2 |
| 344.187 | 4.73 | TTOF0473 (m/z: 344.1870), TTOF0590 (m/z: 344.1871) | 2 |
| 313.1731 | 5.07 | TTOF0768 (m/z: 313.1730), TTOF0827 (m/z: 313.1733) | 2 |
| 445.1603 | 5.23 | TTOF0874 (m/z: 445.1603), TTOF0875 (m/z: 445.1602) | 2 |
| 396.1663 | 6.29 | TTOF0351 (m/z: 396.1663), TTOF0944 (m/z: 396.1664) | 2 |
| 566.2922 | 7.75 | TTOF0881 (m/z: 566.2917), TTOF0917 (m/z: 566.2928) | 2 |
| 548.221 | 8.15 | TTOF0601 (m/z: 548.2209), TTOF0755 (m/z: 548.2211) | 2 |
| 513.2043 | 8.74 | TTOF0539 (m/z: 513.2039), TTOF0579 (m/z: 513.2046) | 2 |
| 445.1459 | 9.27 | TTOF0158 (m/z: 445.1463), TTOF0356 (m/z: 445.1455) | 2 |
| 376.1624 | 9.51 | TTOF0731 (m/z: 376.1626), TTOF0957 (m/z: 376.1621) | 2 |
| 371.1599 | 9.65 | TTOF0774 (m/z: 371.1600), TTOF0926 (m/z: 371.1599) | 2 |
| 397.1741 | 10.26 | TTOF0845 (m/z: 397.1736), TTOF0960 (m/z: 397.1746) | 2 |
| 396.1713 | 10.27 | TTOF0673 (m/z: 396.1717), TTOF0863 (m/z: 396.1710) | 2 |
| 398.1986 | 10.95 | TTOF0184 (m/z: 398.1978), TTOF0392 (m/z: 398.1994) | 2 |
| 313.1737 | 12.32 | TTOF0001 (m/z: 313.1734), TTOF0828 (m/z: 313.1740) | 2 |
| 434.1694 | 12.73 | TTOF0291 (m/z: 434.1688), TTOF0885 (m/z: 434.1701) | 2 |
| 376.1458 | 12.96 | TTOF0004 (m/z: 376.1464), TTOF0179 (m/z: 376.1451), TTOF0575 (m/z: 376.1458) | 3 |
| 337.2172 | 13.04 | TTOF0090 (m/z: 337.2176), TTOF0775 (m/z: 337.2167) | 2 |
| 299.0818 | 13.34 | TTOF0759 (m/z: 299.0814), TTOF0961 (m/z: 299.0823) | 2 |
| 376.1404 | 13.77 | TTOF0818 (m/z: 376.1400), TTOF0918 (m/z: 376.1408) | 2 |
| 299.0793 | 14.19 | TTOF0758 (m/z: 299.0795), TTOF0964 (m/z: 299.0790) | 2 |
| 471.1801 | 14.58 | TTOF0853 (m/z: 471.1799), TTOF0936 (m/z: 471.1803) | 2 |
| 445.1622 | 14.76 | TTOF0379 (m/z: 445.1627), TTOF0385 (m/z: 445.1618) | 2 |
| 285.1036 | 14.96 | TTOF0549 (m/z: 285.1035), TTOF0905 (m/z: 285.1037) | 2 |
| 442.2223 | 17.92 | TTOF0065 (m/z: 442.2221), TTOF0660 (m/z: 442.2224) | 2 |
| 285.1053 | 18.04 | TTOF0506 (m/z: 285.1058), TTOF0915 (m/z: 285.1049) | 2 |
| 533.2272 | 18.3 | TTOF0753 (m/z: 533.2272), TTOF0763 (m/z: 533.2272) | 2 |
| 434.1704 | 18.49 | TTOF0440 (m/z: 434.1710), TTOF0849 (m/z: 434.1698) | 2 |
| 371.1608 | 18.84 | TTOF0025 (m/z: 371.1612), TTOF0599 (m/z: 371.1605) | 2 |
| 365.1968 | 21.05 | TTOF0431 (m/z: 365.1973), TTOF0491 (m/z: 365.1963) | 2 |
| 376.1571 | 22.76 | TTOF0734 (m/z: 376.1569), TTOF0956 (m/z: 376.1572) | 2 |
| 566.291 | 25.4 | TTOF0424 (m/z: 566.2902), TTOF0574 (m/z: 566.2918) | 2 |
| 402.193 | 26.33 | TTOF0279 (m/z: 402.1926), TTOF0569 (m/z: 402.1935) | 2 |
| 400.1427 | 27.69 | TTOF0777 (m/z: 400.1427), TTOF0867 (m/z: 400.1427) | 2 |
| 370.1368 | 28.14 | TTOF0655 (m/z: 370.1364), TTOF0873 (m/z: 370.1372) | 2 |
| 309.1058 | 30.81 | TTOF0638 (m/z: 309.1059), TTOF0757 (m/z: 309.1057) | 2 |
| 313.177 | 32.26 | TTOF0769 (m/z: 313.1768), TTOF0829 (m/z: 313.1772) | 2 |

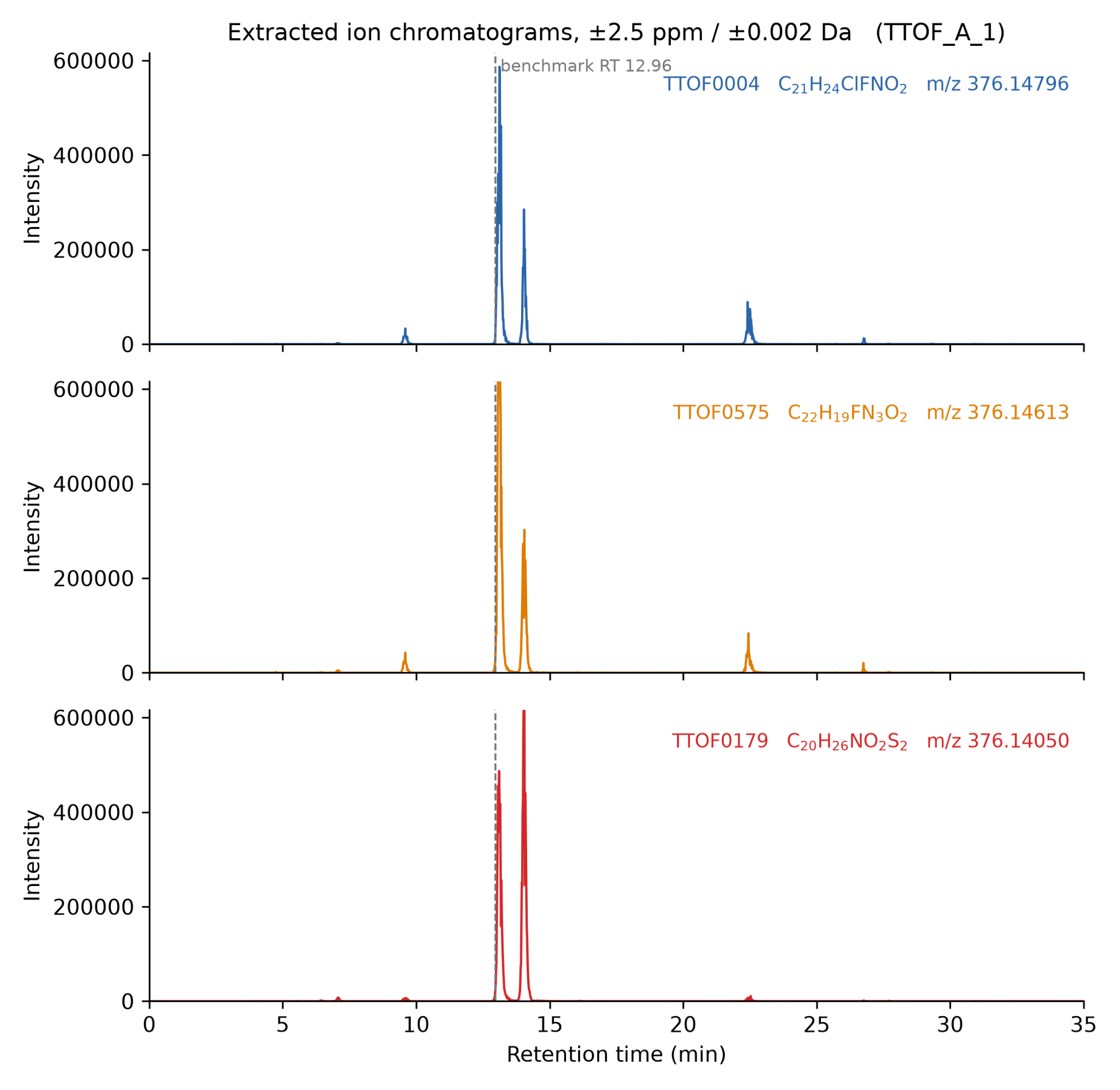

**Supplementary 2 Figure 1.** Extracted ion chromatograms of the three benchmark entries reported at m/z 376.146 and 12.96 min on the SCIEX TripleTOF 6600 (sample TTOF_A_1). Each panel is one entry, extracted at the [M+H]+ monoisotopic mass of its molecular formula, with a tolerance of 2.5 ppm or 0.002 Da, whichever is larger. Panels share both axes. The dashed line marks the retention time given in the benchmark list, 12.96 min.

Three entries at m/z 376.146 (TripleTOF 6600). The ground truth puts TTOF0004, TTOF0575 and TTOF0179 at 12.96 min with almost the same m/z. We took their molecular formulas, calculated the [M+H]+ monoisotopic mass of each with ChemCalc^19^, and extracted an EIC at each mass over the whole run. The three masses are 376.14796, 376.14613 and 376.14050, which are 20 ppm apart, so the compounds are not isobaric. Each EIC shows four peaks, at 9.60, 13.05, 14.03 and 22.50 min. We then measured the m/z at each apex 376.14650 at 13.05 min, matching C22H18FN3O2, and 376.14115 at 14.03 min, matching C20H25NO2S2. TTOF0179 therefore elutes 1.07 min later than the list states, and only two of the three compounds share the peak at 13.05 min. We scored against the published values without correction.
